# Histological, transcriptomic and *in vitro* analysis reveal an intrinsic activated state of myogenic precursors in hyperplasic muscle of trout

**DOI:** 10.1101/332585

**Authors:** Sabrina Jagot, Nathalie Sabin, Aurélie Le Cam, Jérôme Bugeon, Pierre-Yves Rescan, Jean-Charles Gabillard

## Abstract

**Background:** The dramatic increase in myotomal muscle mass in post-hatching fish is related to their ability to lastingly produce new muscle fibres, a process termed hyperplasia. The molecular and cellular mechanisms underlying fish muscle hyperplasia largely remain unknown. In this study, we aimed to characterize intrinsic properties of myogenic cells originating from fish hyperplasic muscle. For this purpose, we compared *in situ* proliferation, *in vitro* cell behavior and transcriptomic profile of myogenic precursors originating from hyperplasic muscle of juvenile trout (JT) and from non-hyperplasic muscle of fasted juvenile trout (FJT) and adult trout (AT).

**Results:** For the first time, we showed that myogenic precursors proliferate in hyperplasic muscle from JT as shown by *in vivo* BrdU labeling. This proliferative rate was very low in AT and FJT muscle. Transcriptiomic analysis revealed that myogenic cells from FJT and AT displayed close expression profiles with only 64 differentially expressed genes (BH corrected p-val < 0.001). In contrast, 2623 differentially expressed genes were found between myogenic cells from JT and from both FJT and AT. Functional categories related to translation, mitochondrial activity, cell cycle, and myogenic differentiation were inferred from genes up regulated in JT compared to AT and FJT myogenic cells. Conversely, Notch signaling pathway, that signs cell quiescence, was inferred from genes down regulated in JT compared to FJT and AT. In line with our transcriptomic data, *in vitro* JT myogenic precursors displayed higher proliferation and differentiation capacities than FJT and AT myogenic precursors.

**Conclusions:** The transcriptomic analysis and examination of cell behavior converge to support the view that myogenic cells extracted from hyperplastic muscle of juvenile trout are intrinsically more potent to form myofibres than myogenic cells extracted from non-hyperplasic muscle. The generation of gene expression profiles in myogenic cell extracted from muscle of juvenile trout may yield insights into the molecular and cellular mechanisms controlling hyperplasia and provides a useful list of potential molecular markers of hyperplasia.

## Background

Post-hatching muscle growth in most teleost fish occurs in two processes. The first process which is common with amniotes refers to increase of fibre size and is termed hypertroph. The second process refers to the formation of new muscle fibers throughout the entire myotome and is termed hyperplasia [1–3]. A persistence of hyperplasic growth after juvenile stage was reported in large final size fish as gilthead bream [4], carp [5], european sea bass [6] and rainbow trout [7, 8]. Nevertheless, this production of new muscle fibers decreases with age [7], and hyperplasia was no longer observed in 18-months old trout [8]. Furthermore, it is well known that fasting stops growth [9] and an inhibition of *in vitro* proliferation of myogenic precursors in fasted rainbow trout has been observed [10].

Muscle hyperplasia requires muscle stem cells, also called satellite cells [11] which are localized between myofibre and basal lamina. Once activated during development, growth or after muscle injury, myogenic precursors proliferate and differentiate to eventually form nascent myofibres [12, 13]. Satellite cells have been clearly identified *in situ* in muscle of carp [14] and zebrafish [15]. *In vitro*, myogenic precursors extracted from trout and carp muscle proliferate and fuse into myotube [10, 16, 17]. Whether myogenic progenitors of fish hyperplasic muscle exhibit specific physiological state is largely unknown. To test this hypothesis, we extracted these cells from hyperplasic muscle of juveniles growing trout (JT), and non-hyperplasic muscle of fasted juvenile trout (FJT) and adult trout (AT), and compared their ability to proliferate *in situ*, their transcriptome and their proliferation and differentiation capacities in culture.

Our results converge to support the view that myogenic cells extracted from hyperplasic muscle of juvenile trout are intrinsically more potent than myogenic cells extracted from non-hyperplasic muscle.

## Results

### Myogenic precursors proliferate in hyperplasic muscle during post-larval growth

In order to quantify the number of proliferative satellite cells in trout of 5g, 500g and fasted trout of 5g, we developed immunofluorescence analysis to spot proliferative nuclei in satellite cell position, i.e. located under the basal lamina. For this purpose, we injected fish with BrdU and performed immunofluorescence analysis with an antibody against BrdU and laminin a major component of basal lamina. As shown in figure 1, the percentage of BrdU positive nuclei in juvenile trout was 7,2%, whereas this proportion dropped to 1.3% in larger trout (500g) and 0,1% in 3-week fasted juvenile trout.

**Figure 1:**
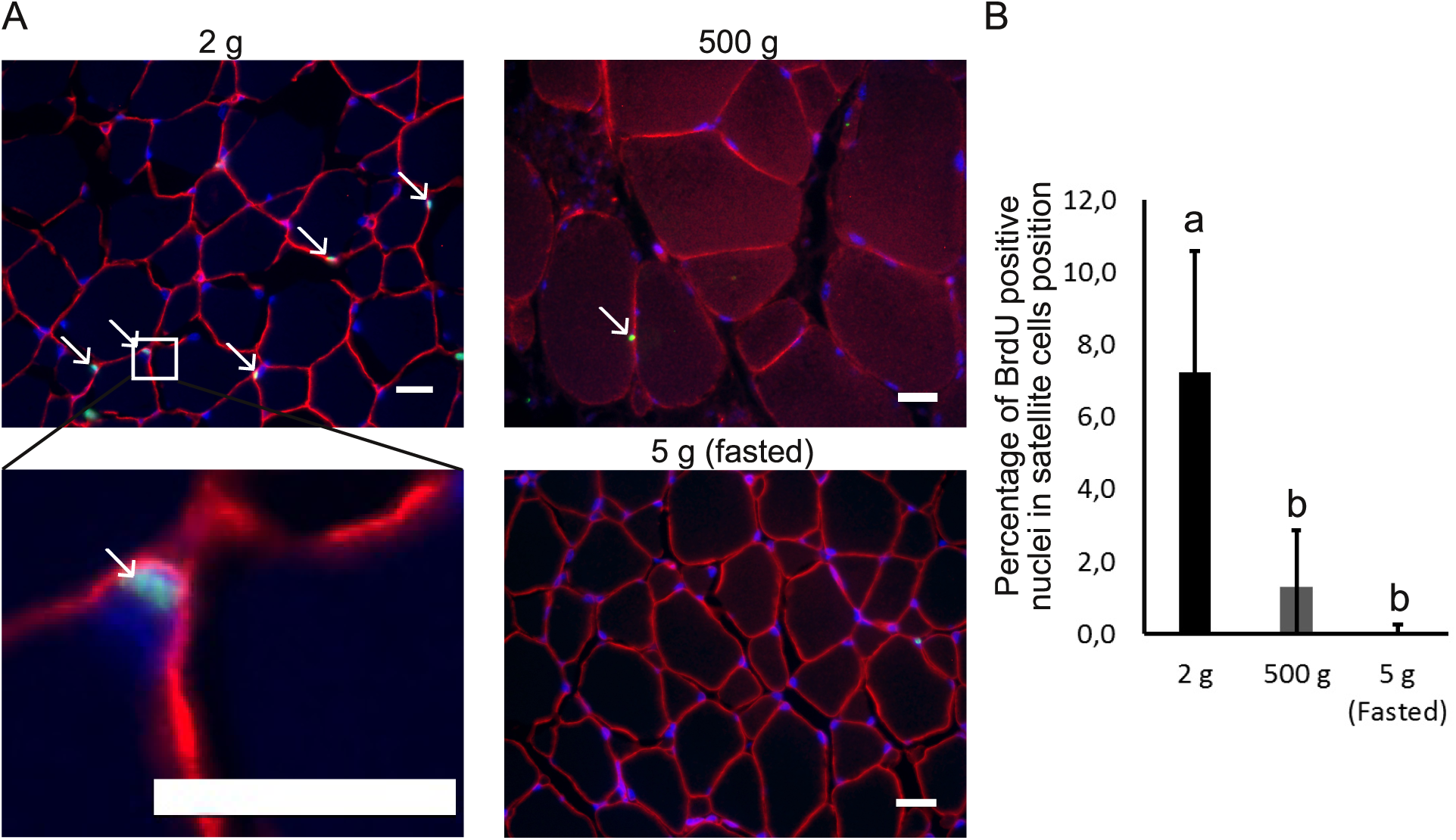
Quantification of satellite cells proliferation in hyperplasic and non-hyperplasic muscle of trout. (A) Muscle cross sections stained with anti-laminin (red) and anti-BrdU (green) in trout of 2g, 500g and of 3-weeks fasted trout (5g). Nuclei were counter-stained with DAPI (blue) (scale bar = 20µm). (B) Quantification of BrdU positive nuclei (% ±SD) in satellite cells position, (under the basal lamina), in white muscle of trout weighing 2g, 500g and of 3-weeks fasted trout weighing 5g. Different letters indicates a significant difference between means (Kruskal-Wallis and Dunn’s multiple comparisons test; p-value ≤ 0.05; n = 5).

### Myogenic precursors extracted from hyperplasic and non-hyperplasic trout muscles exhibit distinct transcriptome

To better known the intrinsic molecular properties of myogenic precursors from hyperplasic muscle, we compared the transcriptome of myogenic precursors extracted from juvenile trout (JT) displaying hyperplasic muscle growth with that of myogenic precursors extracted from non-hyperplasic muscle resulting from fasted juvenile trout (FJT) and adult trout (AT). For this purpose, we first compared gene expression profiles between myogenic precursors from FJT and AT in order to identify genes whose differential expression would be specifically related to age or fasting. LIMMA statistical test [18] (BH corrected p-val < 0.001) showed that only 64 genes were differentially expressed between FJT and AT samples. These differentially expressed genes (DEGs) were subsequently discarded for further analysis. Using two LIMMA statistical tests (BH corrected p-val < 0.001) a total of 3992 DEGs were identified between JT and FJT and 4253 DEGs between JT and AT. Then, we retained common genes found in this two differential analysis and found a total of 2623 differentially expressed genes between hyperplasic (JT) and non-hyperplasic muscle (FJT and AT). These differentially expressed genes were then hierarchically clustered. The unsupervised clustering, which is shown in figure 2 (available as supplemental data file 1), resulted in the formation of two major gene clusters. The cluster 1 comprised 1865 genes up regulated in JT myogenic precursors and cluster 2 comprised 758 genes down regulated in JT myogenic precursors.

**Figure 2:**
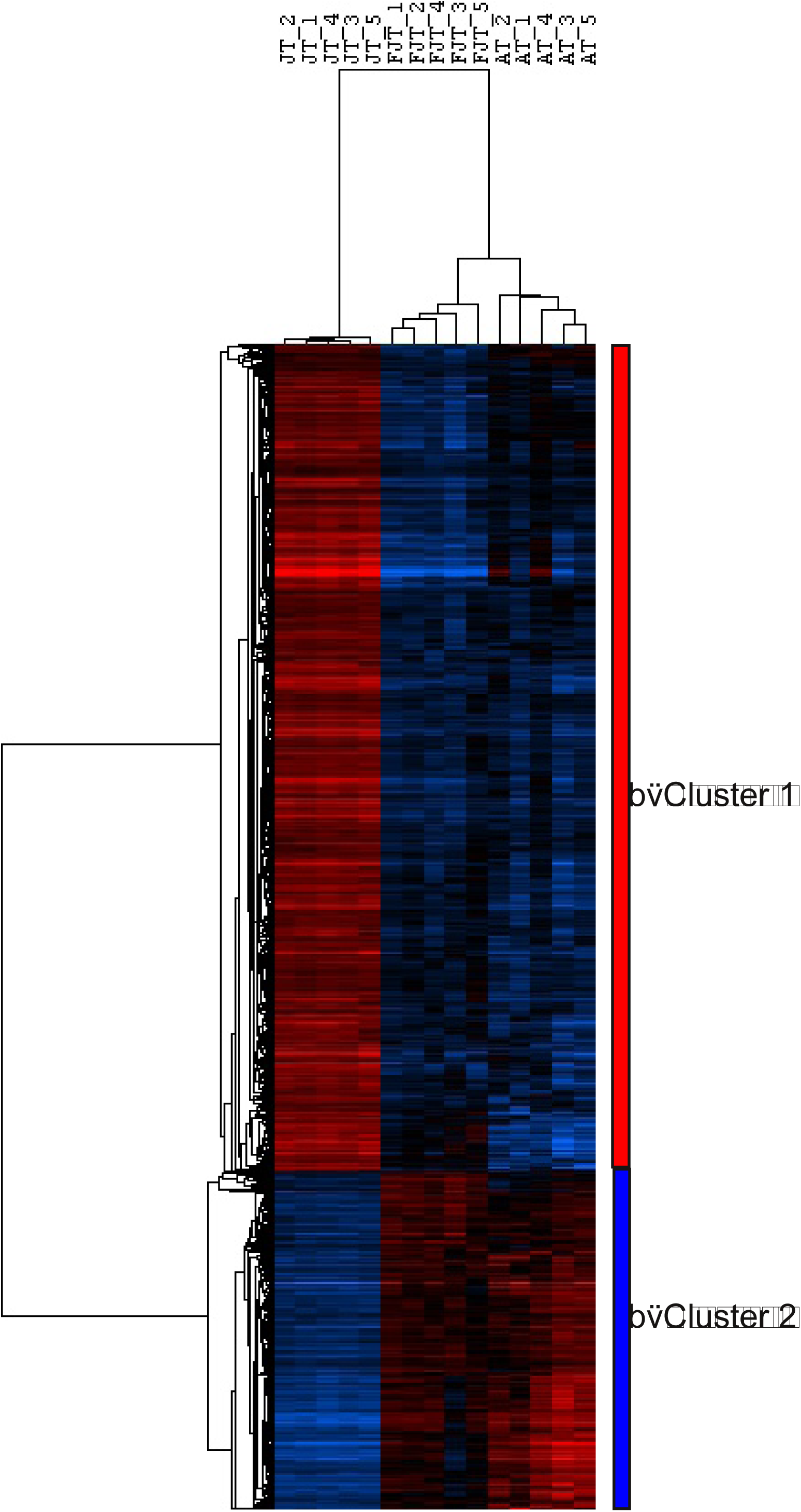
Hierarchical clustering of differentially expressed genes between JT myogenic precursors and FJT and AT myogenic precursors. Each row represents the expression pattern of a single gene and each column corresponds to a single sample: columns 1 to 5: JT myogenic precursors sampled; columns 6 to 10: FJT myogenic precursors sampled; and columns 11 to 15: AT myogenic precursors sampled. The expression levels are represented by colored tags, with red representing the highest levels of expression and blue representing the lowest levels of expression.

### JT myogenic precursors exhibit transcriptomic signature of activated state cell

DAVID analysis of the 1206 eligible genes from cluster 1 revealed significant enrichment (table 1) in genes involved in translation (p=2.8E^−26^), mitochondrial activity (p=3,85E^−11^) and oxidative phosphorylation (p=7,31E^−12^). Among other significant functional categories inferred from up regulated genes in JT myogenic precursors, we found the GO term mitotic cell cycle (p=2.26E^−20^). Genes belonging to this functional category included genes encoding cell division cycle (cdc) proteins (8), cyclin dependent kinases (6), cyclins (6), genes involved in chromosomes segregation (20) as shown in figure 3. Enrichment in gene involved in DNA metabolic process and replication such as minichromosome maintenance complex components, non-homologous end-joining factor1, DNA polymerases, DNA primases, DNA topoisomerases, replication proteins were also found. Cluster 1 also included many genes encoding epigenetic transcriptional regulators. Among them were Swi/Snf chromatin enzymes and several DNA (cytosine-5-)-methyltransferases. We found also many genes encoding extracellular components including collagens (14 genes), laminin subunits (3 genes) and entactin as well as genes contributing to the formation of the myofibrils (i.e, 8 genes encoding myosins, 5 genes encoding troponins and 3 genes encoding tropomyosins). At last, besides the large number of myofibrillary proteins, we found many genes involved in myoblast differentiation and fusion such as *six1b, six4b, mef2d, myogenin, tmem8c* (*myomaker*), *muscle creatine kinase* (figure 4). Overall, cluster 1 showed enrichment in genes involved in protein synthesis, cell division and myogenic differentiation.

**Table 1:** **Functional categories inferred from up regulated genes in JT myogenic precursors**. Table of the most significant Gene Ontology terms in Biological Process and Cellular Component that were found following functional enrichment analysis (DAVID Software 6.7) among genes up regulated in JT myogenic precursors.

| GO terms Biological Process | Number of genes | p-value |
| --- | --- | --- |
| GO:0006412 translation | 84 | 2,82E-26 |
| GO:0006119 oxidative phosphorylation | 30 | 7,31E-12 |
| GO:0042775 mitochondrial ATP synthesis coupled electron transport | 22 | 3,85E-11 |
| GO:0000278 mitotic cell cycle | 80 | 2,26E-20 |
| GO:0000280 nuclear division | 51 | 2,38E-14 |
| GO:0048285 organelle fission | 52 | 3,10E-14 |
| GO:0007059 chromosome segregation | 20 | 1,65E-06 |
| GO:0006260 DNA replication | 30 | 3,83E-05 |
| GO:0006259 DNA metabolic process | 61 | 2,01E-05 |
| GO terms Cellular Component | Number of genes | p-value |
| GO:0005840 ribosome | 72 | 1,61E-29 |
| GO:0030529 ribonucleoprotein complex | 102 | 1,59E-21 |
| GO:0005739 mitochondrion | 211 | 1,77E-44 |
| GO:0070469 respiratory chain | 29 | 6,83E-14 |
| GO:0005839 proteasome core complex | 10 | 3,84E-06 |
| GO:0000776 kinetochore | 16 | 3,10E-04 |
| GO:0030017 sarcomere | 13 | 4,64E-02 |

**Figure 3:**
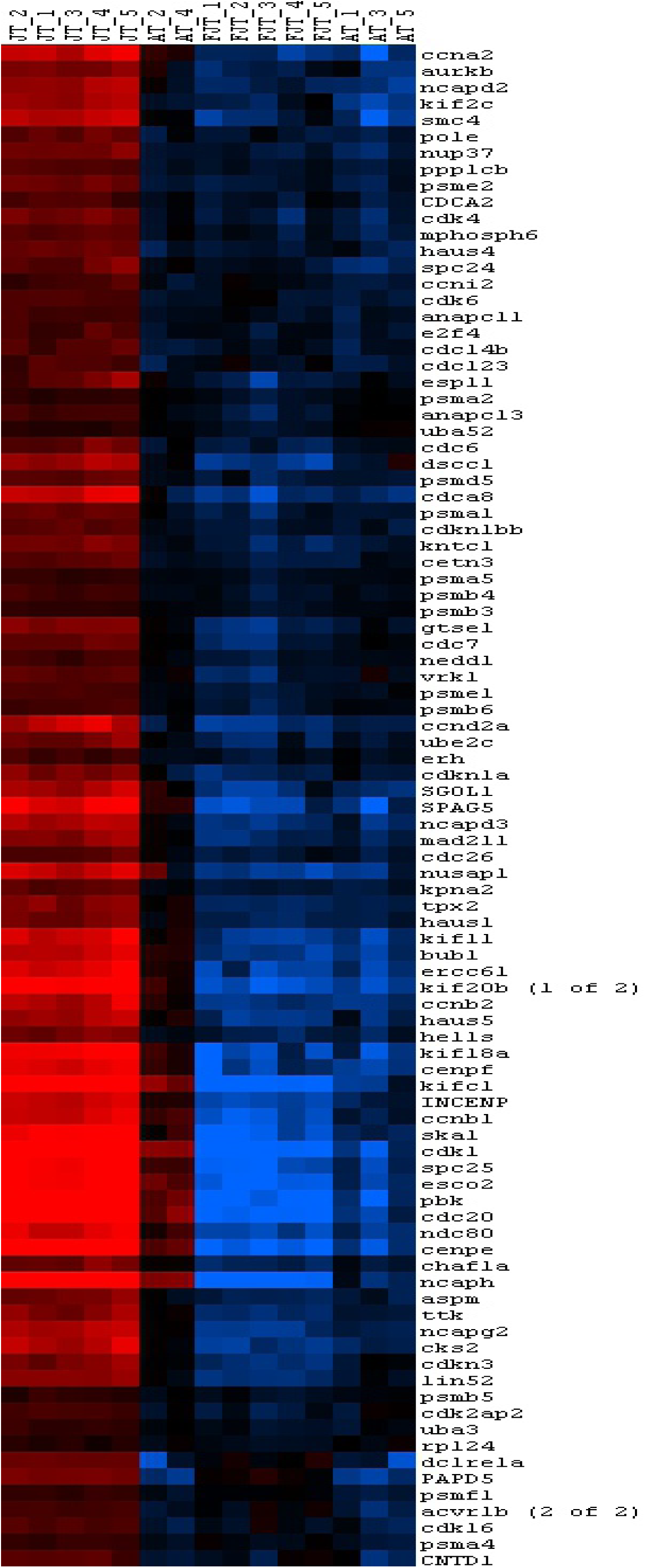
Hierarchical clustering of differentially expressed cell cycle genes between JT myogenic precursors and FJT and AT myogenic precursors. Each row represents the expression pattern of a single gene and each column corresponds to a single sample: columns 1 to 5: JT myogenic precursors sampled; columns 6 to 10: FJT myogenic precursors sampled; and columns 11 to 15: AT myogenic precursors sampled. The expression levels are represented by colored tags, with red representing the highest levels of expression and blue representing the lowest levels of expression.

**Figure 4:**
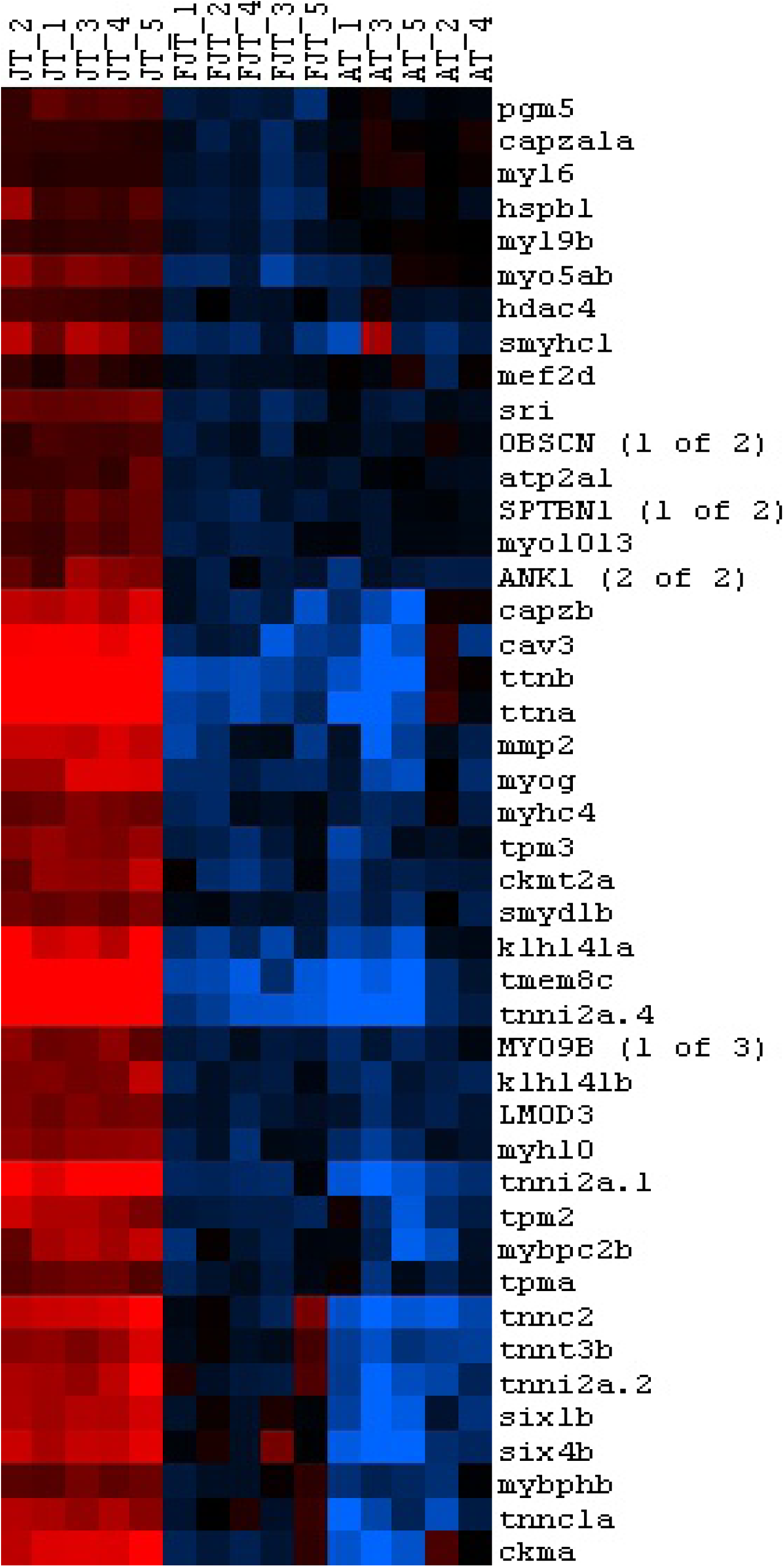
Hierarchical clustering of differentially expressed myogenic genes between JT myogenic precursors and FJT and AT myogenic precursors. Each row represents the expression pattern of a single gene and each column corresponds to a single sample: columns 1 to 5: JT myogenic precursors sampled; columns 6 to 10: FJT myogenic precursors sampled; and columns 11 to 15: AT myogenic precursors sampled. The expression levels are represented by colored tags, with red representing the highest levels of expression and blue representing the lowest levels of expression.

### Genes associated with the quiescent state are down regulated in JT myogenic precursors

Cluster 2 comprised genes that were down regulated in JT myogenic precursors compared to both FJT and AT myogenic precursors. In this cluster, we identified genes of the Notch pathway, suggesting a repression of quiescent state. Associated with this quiescency state pathway we found *jagged1b, jagged2b, dll4, dlc, notch1a, notch1b, notchl, her6* and *hey1* among genes contained in cluster 2. We detected some genes which play repression roles in proliferation as *hexim1b* [19], *stat3* [20], and *Dach1* also known to inhibit Six protein activity [21]. Among the down regulated genes in JT myogenic precursors, we distinguished genes which plays repression roles in myogenic differentiation as *ddit3* [22], *trim33* [23], *bhlhe40* [24], *tal1* [25]. Moreover, a marker of quiescent satellite cells [26], *nestin* was down regulation in JT myogenic precursors. We also observed a global repression of the TGFβ pathway in JT myogenic precursors. Indeed, 7 genes involved in TGFβ pathway, were down regulated in JT myogenic precursors (*tgfb2, tgfbr1, bmpr2b, bmpr1bb, smad3b, smad6a* and *acvrrl1*) whereas 5 inhibitors of TGFβ pathway were up regulated in JT myogenic precursors (*Bmp7a, gremlin2, dcn, fstl1b* and *fsta*). Overall, cluster 2 showed enrichment in genes involved in inhibition of proliferation, repression of myogenic differentiation and maintenance of cellular quiescent state.

### JT myogenic precursors have an enhanced intrinsic capacity of for *in vitro* proliferation

To know more about the intrinsic molecular properties of myogenic precursors of hyperplasic muscle compared to myogenic precursors of non-hyperplasic muscle, we carried out a primary cell culture of myogenic progenitors extracted from JT, FJT and AT conditions. Cell proliferation assays using BrdU showed a higher proliferation rate of JT myogenic precursors (40.1%) after two days of culture compared to FJT (0.8%) and AT (10.3%) myogenic precursors (figure 5). Then, to determine whether the transcriptomic activation signatures were related to a differential cell behavior regarding proliferating capacity, we also measured the proliferation rate of JT, FJT and AT myogenic cells at 5, 8 and 11 days after plating. In JT myogenic precursors the proliferation rate increased from D2 to reach a maximum rate at D5 with 61.4% of BrdU positive nuclei, then the proliferation rate decreased from D8 to 42.4% to D11 to 31.4%. In sharp contrast, proliferation rate of FJT myogenic precursors remained low and tended to increase up to 12,9% at D8. For AT myogenic precursors, the proliferation rate increased at D5 to 48.3% to almost reach the proliferation rate in JT myogenic precursors and decreased from D8 to 31.6% and at D11 to 19.1%. Thus, the kinetic of proliferation of the AT precursors was close to that one of JT but with a lower rate from D5 to D11. Overall, myogenic precursors of JT exhibit a global enhanced proliferation capacity under *in vitro* conditions compared to FJT and AT.

**Figure 5:**
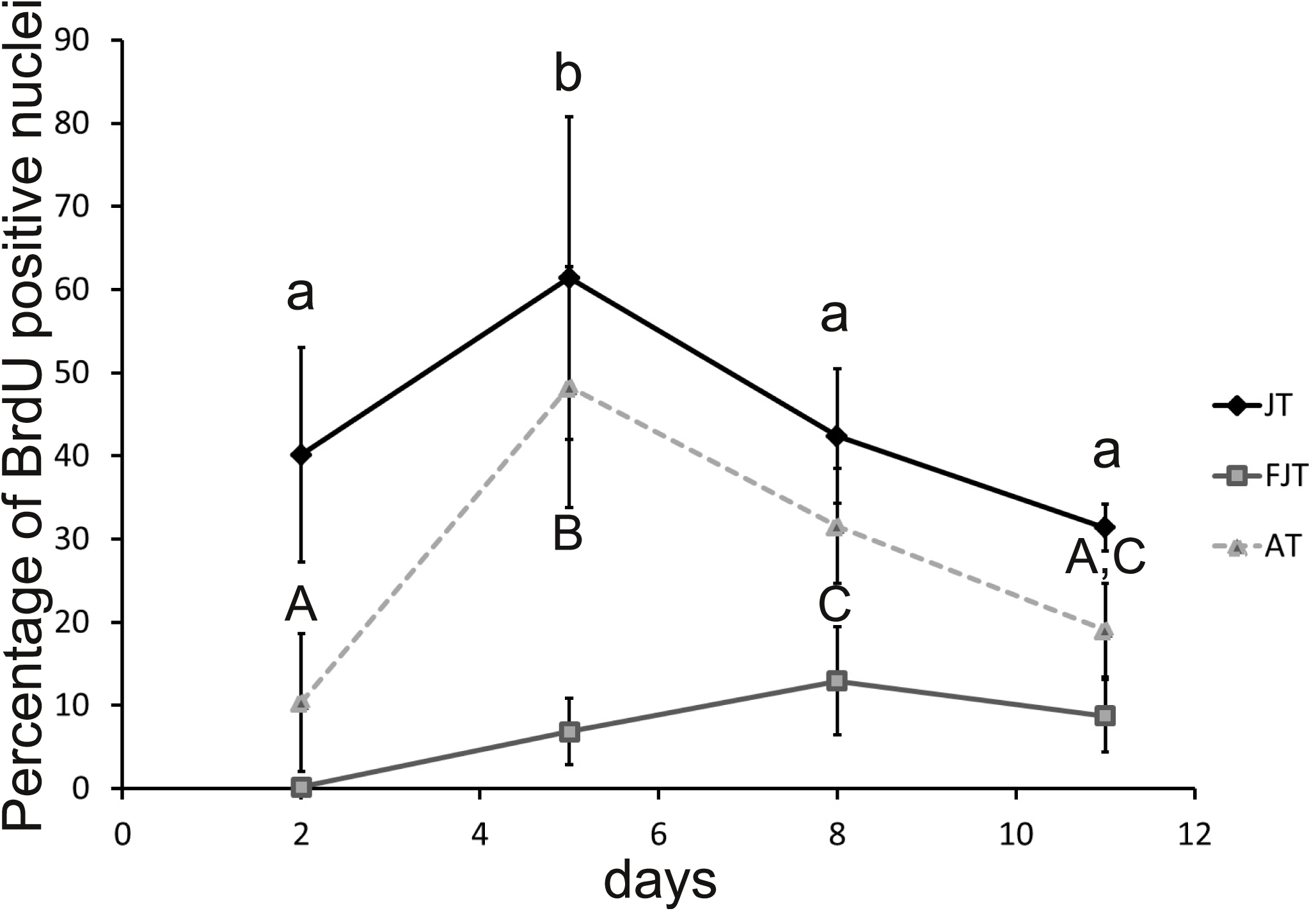
Proliferation rate of JT, FJT and AT myogenic precursors after 2, 5, 8, 11 days of plating (D2, D5, D8 and D11). Each point represents the mean (% ±SD) of BrdU positive nuclei ratio for each condition at D2, D5, D8 and D11. Different letters indicates a significant difference between means (two-way ANOVA and Tukey’s multiple comparisons test; p-value ≤ 0.05; n ≥ 5).

### JT myogenic precursors have an enhanced capacity for *in vitro* myogenic differentiation

To go further on the characterization of the intrinsic molecular properties of myogenic precursors of hyperplastic muscle, we quantified the *in vitro* differentiation capacities of JT, FJT and AT myogenic precursors. At D2, we observed an extremely low differentiation rate in JT (1.4%), FJT (1%) and AT (1.6%) myogenic precursors (figure 6). This result indicates that very few myocytes were extracted at the beginning of the cell culture. Then, we also measured the differentiation rate at 5, 8 and 11 days after plating of JT, FJT and AT myogenic progenitors. In JT myogenic precursors the differentiation rate increased at D5 to 11.6%, D8 to 24.4% and reach a maximum rate at D11 with 28% of nuclei contained in myosin positive cells. In sharp contrast, the differentiation of FJT precursors remained very low during the first 8 days (<0.5%) then a differentiation resumption was observed at D11 (6.4%). For AT myogenic precursors, no significant increase of the differentiation rate was observed even after 11 days of culture. Overall, JT myogenic precursors exhibited a global enhanced differentiation capacity under *in vitro* conditions.

**Figure 6:**
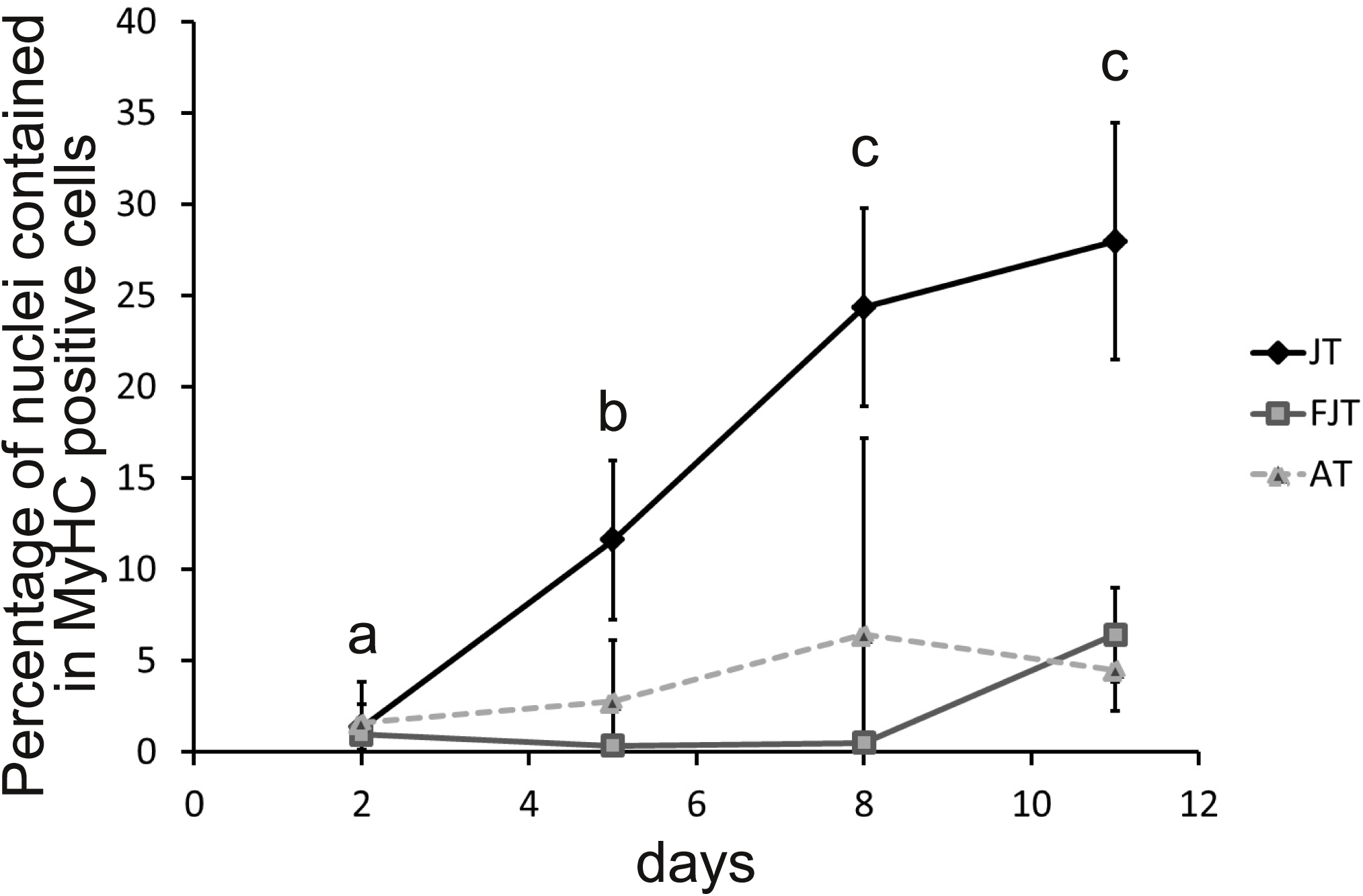
Differentiation rate of JT, FJT and AT myogenic precursors after 2, 5, 8, 11 days in culture (D2, D5, D8 and D11). Each point represents the mean (% ±SD) of the percentage of nuclei contained in MyHC positive cells for each condition at D2, D5, D8 and D11. Different letters indicates a significant difference between means (two-way ANOVA and Tukey’s multiple comparisons test; p-value ≤ 0.05; n ≥ 6).

Evaluation of the expression level by qPCR of *myogenin* and *myomaker* after 2 days in cell culture validated the transcriptomic results as shown in figure 7. Indeed, the expression of *myogenin* and *myomaker* were higher in JT myogenic precursors compared to AT and FJT myogenic precursors. In addition, the expression level of *myogenin* and *myomaker* after 8 days in culture increase in FJT myogenic precursors. These were contrasting with expression level in AT myogenic precursor that did not exhibit such an increase between D2 and D8. Overall, qPCR data validated our previous results with JT myogenic precursors as more engaged in differentiation program than AT and FJT myogenic precursors.

**Figure 7:**
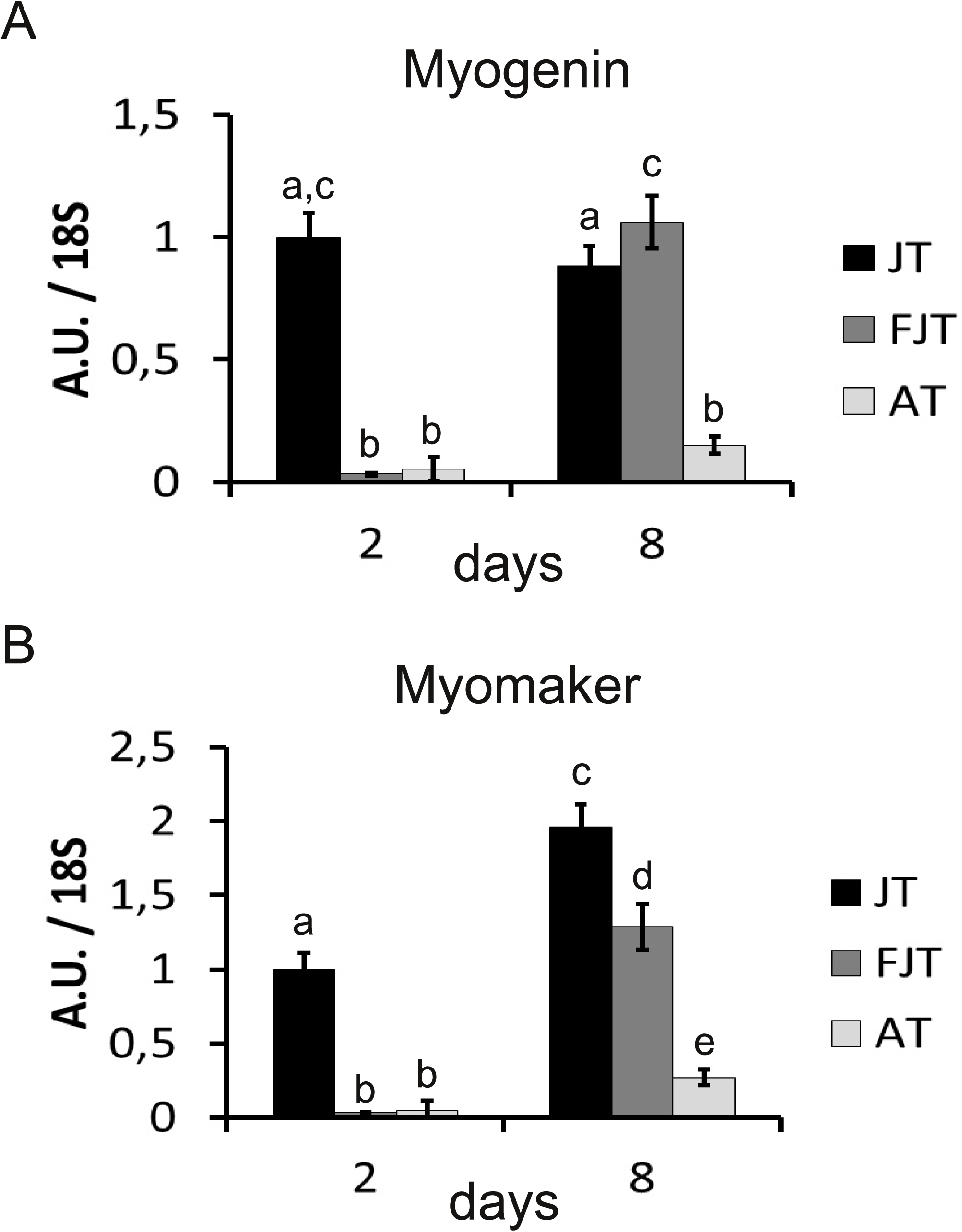
Quantification of the expression of *myogenin* and *myomaker* in JT, FJT and AT myogenic precursors. Each bar represents the mean (AU ±SD) of the expression of *myogenin* (A) and *myomaker* (B) normalized by the expression mean of 18S as referential gene for each condition at D2 and D8. Different letters indicates a significant difference between means (two-way ANOVA and Tukey’s multiple comparisons test; p-value ≤ 0.05; n ≥ 4).

## Discussion

Post-hatching muscle growth in most teleost such as trout, lastingly occurs by fiber hypertrophy and formation of new muscle fibers. This latter process, termed hyperplasia, requires proliferation, differentiation and fusion of muscle stem cells (satellite cells) to form new multinucleated myofibers. We examined in this study the hypothesis that post-hatching muscle hyperplasia in fish is associated with a peculiar physiological status of myogenic precursors predetermining them to self-renew and differentiate. For this purpose, we examined proliferation of trout satellite cells *in vivo* and compared gene expression profiling and *in vitro* myogenic potential of satellite cells extracted from juvenile trout muscle displaying intense hyperplastic growth (JT), with satellite cells extracted from trout muscle that no longer exhibited muscle hyperplasia, namely fasted juvenile trout (FJT) and adult trout (AT).

Many studies on mammalian isolated satellite cells were carried out on cells directly isolated from muscle and purified by FACS using fluorescent reporters or cell surface marker [27]. As these technologies cannot yet be used in trout fish, we took advantage of the specific adhesion of satellite cells on laminin substrate to enrich them in culture [17, 28]. Although it has been reported that isolation procedures alter gene expression of myogenic precursors [29, 30], we assumed in this study that the differential *ex vivo* properties of trout satellite cells originating either from hyperplastic or non-hyperplastic muscle, somehow reflect intrinsic differences preexisting before their extraction from muscle.

First, we sought to identify and quantify proliferative satellite cells in muscle of growing *versus* non-growing trout using *in vivo* BrdU injection followed by double immuno-labeling of laminin and BrdU. In agreement with Alfei et al (1989)[31], our results clearly evidenced a higher rate of BrdU positive cells in muscle of JT compared to FJT and AT, notably at sites corresponding to the satellite cell niche. This shows that fish hyperplastic muscle contains proliferative satellite cells well after hatching, what sharply contrasts with the mitotic quiescence of satellite cells located in mature mouse muscle [32].

Relative to satellite cells from non-hyperplastic muscle, satellite cells from juvenile trout were found to exhibit up-regulated gene set related to high metabolic activity as shown by enrichment in genes involved in translational efficiency and genes encoding structural and functional components of mitochondria, notably those involved in energy production for execution of biosynthesis events. Mitochondrial biogenesis has been associated with the shift from quiescence to proliferation of satellite cells [33, 34]. In keeping with this, our result that matches meta-analyses of multiple transcriptomes revealing low expression of genes associated with oxidative phosphorylation in mouse quiescent satellite cells [35], supports the view that JT cells are intrinsically activated compared to satellite cells from non-hyperplastic muscle. Other major functional categories inferred from genes up-regulated in myogenic precursors derived from hyperplastic muscle were related to DNA replication and cell cycle. This finding, which is quite in agreement with the proliferation rate of these cells measured *in vivo* and *ex vivo*, strongly reinforces the view that satellite cells isolated from trout hyperplastic muscle are in an activated state. Also, several major genes signing myogenic differentiation were found to be overexpressed. Among them were *myogenin* which invalidation prevents myogenic differentiation in mouse [36] and *myomaker* which is necessary for myoblast fusion into myotube as shown by gene invalidation [37]. In keeping with this, it is interesting to note that mitochondrial activity, which is higher in JT satellite cells relative to FJT and AT cells, has been reported to positively regulate myogenesis [38]. Conversely, transcriptome of FJT and AT myogenic precursors, compared to that of JT myogenic precursors, revealed up regulation of genes involved in maintenance of stem cell quiescence, notably genes involved in Notch signaling [39] or known as marker of quiescent muscle stem cell. These results are in agreement with data obtained in mouse showing an up regulation of *notch* and *Hey* genes in quiescent satellite cells [40]. In addition, the up regulation of several genes involved in TGFbeta pathway was in line with a repression of differentiation of myogenic precursors [41]. Indeed, we notably observed an up-regulation of BMP receptor type 1 which knock-down in mouse satellite cells caused premature myogenic differentiation [42]. All these data support the view that satellite cells extracted from muscle of fasted trout or adult trout are close to a quiescent state compared to satellite cells from juvenile trout.

Another major result of our study was that behavior of satellite cells from hyperplastic muscle quite differs from that of satellite cells extracted from non-hyperplastic muscle. Specifically, we found that cultured JT myogenic precursors exhibited higher proliferation rate and differentiation capacities than FJT and AT myogenic precursors. These observations, that match transcriptome data, further support the view that myogenic cells from hyperplastic muscle of juvenile trout are intrinsically more potent to form myofibres than satellite cells from non-hyperplastic muscle.

What could determine intrinsic myogenic capacity of JT cells ? One possible cause, inferred from transcriptome analysis, could relate to epigenetic regulations of transcription. Indeed, up-regulation of genes involved in DNA methylation was found in JT myogenic precursors, notably several DNA methyl transferase (dnmt1, 3ab and 3b) known to be involved in muscle stem cell activation [43]. Furthermore, as previously reported in hyperplastic growth zone of trout larvae [44] and in activated satellite cells of mouse and trout regenerating muscle [45], we observed in JT cells the overexpression of many SWI/SNF chromatin remodeling enzymes, which dynamic recruitment regulate many stages of myogenesis [46].

## Conclusion

The satellite cells from muscle of trout juveniles exhibit *in vivo* and *ex vivo* features of activation that are not found in satellite cells isolated from non-hyperplastic muscle. Thus, muscle hyperplastic growth in fish likely relates to the fact that satellite cells in these animals are intrinsically potent to form myofibres well after hatching.

## Methods

### Animals

Rainbow trout (*Oncorhynchus mykiss*) weighting from 2g to 2kg were raised to a 12 h light:12 h dark photoperiod and 12 ± 1 °C in a recirculating rearing system located in the Laboratory of Physiology and Genomics of Fish. Fish were fed daily *ad libitum* on a commercial diet or starved during 3 or 4 weeks.

### Measurement of satellite cells proliferation in situ

Intra-peritoneal injections (150µg/g of body weight) of BrdU (Roche, no. 280 879), dissolved in a solution composed with NaOH (0.02N) diluted with NaCl 0.9%, were performed on juvenile rainbow trout (*Oncorhynchus mykiss*) (2g, n = 5), 4 weeks fasted juvenile rainbow trout (5g, n = 5) and 400-500g rainbow trout (n = 6) which exhibited a diminution of hyperplasia.

Muscle tissues were fixed in Carnoy fixative solution for 48 h at 4°C, progressively dehydrated and embedded in paraffin. Transverse paraffin sections (10 μm thick) were stained with laminin antibody (DSHB, D18-c) and BrdU labeling and detection kit (Roche Diagnostics, no. 11 296 736 001) was used following the recommendations of manufacturer to measure the proliferation of the cells. Briefly, tissues were incubated for 30 min at 37 °C with mouse IgG1 anti-BrdU (kit: 11296736001, Sigma) and, after 1h incubation at room temperature in saturation buffer (BSA 1%, 04-100-811C in PBST 0.1%), tissues were incubated overnight at 4°C with mouse IgG2a anti-laminin (DSHB, D18-c). The secondary antibody were diluted (1/1000, Alexa 488 anti-IgG1 mouse A21121 to detect BrdU and Alexa 594 anti-IgG2a mouse A21135 to detect laminin) in PBST and applied for 1 h at room temperature. Tissues were then mounted in Mowiol containing 0.5 μg/ml of DAPI. Tissues cross sections were photographed using a Nikon digital camera coupled to a Nikon Eclipse 90i microscope. At least five images were taken per tissues and the number of nuclei BrdU positive localized between basal lamina and myofiber on the total number of nuclei under basal lamina (myo-nuclei) were calculated using cell counter plugin in Fiji software.

### Isolation of trout precursor myogenic cells

For all studies, myogenic precursors were isolated from juvenile trout (5g, JT), from 3-4 weeks fasted juvenile rainbow trout (5g, FJT) and from adult rainbow trout (1.5-2kg, AT) as previously described [17]. Isolated myogenic precursors were plated on poly-L-lysine and laminin-coated plates at 80,000 cells per cm^2^ for every analysis except to proliferation measurement which were 60,000 cells per cm^2^.

### Gene expression analysis

Using TRIzol reagent (Invitrogen, Carlsbad, CA, USA), total RNA were extracted from cells according to the manufacturer’s recommendations. The total RNA (200ng) were reverse transcribed into cDNA using the High Capacity cDNA Reverse Transcription kit, (Applied Biosystems) and random primers, according to the manufacturer’s instructions. Target gene expression levels were determined by qPCR using specific primers (forward primer sequences; *myogenin* : AGCAGGAGAACGACCAGGGAAC, *myomaker* : AATCACTGTCAAATGGTTACAGA, and reverse primer sequences; *myogenin* : GTGTTGCTCCACTCTGGGCTG, *myomaker* : GTAGTCCCACTCCTCGAAGT). Primers were design on two exons to avoid genomic amplification. Quantitative PCR was performed on a StepOnePlus thermocycler (Applied Biosystems) using SYBR FAST qPCR Master Mix (PowerUp SYBR Green Master Mix kit, A25742, Applied Biosystems). Relative quantification of the target gene transcripts was made using 18S gene expression as reference. Quantitative PCR was performed using 10 μl of the diluted cDNA mixed with 300nM of each primer in a final volume of 20 μl. The PCR protocol was initiated at 95°C for 3 min for initial denaturation followed by the amplification steps (20 sec at 95°C followed by 30 sec at 60°C) repeated 40 times. Melting curves were systematically monitored at the end of the last amplification cycle to confirm the specificity of the amplification reaction. Each PCR run included replicate samples (duplicate of PCR amplification) and negative controls (RNA-free samples, NTC).

### Microarray slides

An Agilent-based microarray platform with 8 ×; 60K probes per slide was used (GEO platform record: GPL24910). Microarray data sets have been submitted to the GEO-NCBI with the accession number: GSE113758.

### RNA labeling and hybridization

RNA from (i) five distinct pools of 24H-cultured myogenic precursors from juvenile trout (JT), (ii) five distinct pools of 24H-cultured myogenic precursors from 3-4 weeks fasted juvenile trout (FJT) and (iii) six distinct pools of 24H-cultured myogenic precursors from adult trout (AT) were used for labelling and hybridization. For each sample, 150ng of RNA was Cy3-labelled according to the manufacturer’s instructions (One-Color Microarray-Based Gene Expression Analysis (Low Input Quick Amp Labeling) Agilent protocol). Briefly, RNA was first reverse transcribed, using a polydT-T7 primer, Cy3 was incorporated by a T7 polymerase-mediated transcription and excess dye was washed using an RNeasy kit (Quiagen). The level of dye incorporation was evaluated using a spectrophotometer (Nanodrop ND1000, LabTech). 600 ng of labelled cRNA was then fragmented in the appropriate buffer (Agilent) for 30 minutes at 60°C before dilution (v/v) in hybridization buffer. Hybridizations were performed in a microarray hybridization oven (Agilent) for 17h at 65°C, using two Agilent 8 ×; 60K high-density oligonucleotide microarray slides. Following hybridization, the slides were rinsed in gene expression wash buffers 1 and 2 (Agilent).

### Data acquisition and analysis

Hybridized slides were scanned at a 3-μm resolution using the Agilent DNA microarray Scanner. Data were extracted using the standard procedures contained in the Agilent Feature Extraction (FE) software version 10.7.3.1. One AT sample that did not give good quality signal on microarray was discarded from the gene expression analysis. Arrays were normalized using GeneSpring software version 14.5. Using R software (3.2.2) a LIMMA (3.26.9) statistical test [18] (BH corrected p-val < 0.001) was used to find differentially expressed genes between FJT and AT. Secondly, two LIMMA statistical tests (BH corrected p-val < 0.001) were used to find differentially expressed genes between JT and FJT, and between JT and AT. We kept significant differentially expressed genes with an expression mean in at least one condition above or equal to 6, corresponding at 3 times background (normalized values). Thirdly, we kept commons genes found in this two differential analysis in the same regulation way with JT as referential condition. For clustering analysis, log transformed values were median-centred and an average linkage clustering was carried out using CLUSTER 3.0 software and the results were visualized with TreeView software. GO enrichment analysis was performed using Database for Annotation, Visualization and Integrated Discovery (DAVID 6.7) software tools.

### Analysis of cell proliferation

Cells were cultured in presence of 10µM BrdU during 24H and cells were collected at days 2, 5, 8 and 11. The cells were fixed with ethanol/glycine buffer (100% ethanol, 50 mM glycine, pH 2). A BrdU labeling and detection kit (11296736001, Sigma) was used following the recommendations of manufacturer to measure the proliferation of the cells. Briefly, the cells were incubated for 30 min at 37 °C with mouse anti-BrdU, washed, and then incubated with the secondary antibody anti-mouse FITC for 30 min. Cells were then mounted in Mowiol containing 0.5 μg/ml DAPI. Cells were photographed using a Nikon digital camera coupled to a Nikon Eclipse 90i microscope. Seven images were taken per well and the number of BrdU positive nuclei on the total number of nuclei was automatically calculated using a macro command on Visilog (6.7) software.

### Analysis of cell differentiation

On days 2, 5, 8 and 11 of culture, cells on glass coverslips were briefly washed twice with phosphate-buffered saline (PBS) and fixed for 30 min with 4% paraformaldehyde in PBS. After three washes, cells were saturated for 1 h with 3% BSA, 0.1% Tween-20 in PBS (PBST). Cells were incubated at room temperature for 3 h with the primary antibody anti-myosin heavy chain (MyHC, DSHB, MF20-c) in blocking buffer [17]. The secondary antibody were diluted (1/1000, Alexa 488 A11001) in PBST and applied for 1 h at room temperature. Cells were mounted with Mowiol containing DAPI (0.5 μg/ml). Cells were photographed using a Nikon digital camera coupled to a Nikon Eclipse 90i microscope. Five images were taken per well and the number of nuclei contained in MyHC positive cells on the total number of nuclei was automatically calculated using a macro command on Visilog (6.7) software.

### Statistical analysis

A two-way ANOVA analysis with a Tukey’s *post hoc* multiple comparisons test was performed on qPCR data, proliferation ratio and differentiation ratio. A Kruskal-Wallis test with a Dunn’s *post hoc* multiple comparisons test was performed on *in situ* satellite cells proliferation data. A p-value below 0,05 was considered significant.

## Declarations

### Ethics approval

Fish used in this study were reared and handled in strict accordance with French and European policies and guidelines of the Institutional Animal Care and Use Committee (no. 3312-20 15121511 022362 and 3313-20 15121511 094929), which approved this study.

### Availability of data and material

Gene expression data supporting the results of this article are available in the Gene Expression Omnibus (GEO) repository under the accession number: GSE113758.

### Competing interests

The authors declare that they have no competing interests.

### Funding

This research were funded by National Institute of Agronomic Research (INRA).

## Authors’ contributions

JCG conceived and supervised the study. SJ, AL and NS performed the experiments. SJ, PYR and JCG analysed the data. JB helped for cell proliferation and differentiation quantification. SJ, PYR and JCG wrote the paper. All authors read and approved the final manuscript.

## Acknowledgements

The fellowship of Sabrina Jagot was supported by INRA PHASE and the Région Bretagne. We also thank C. Duret for husbandry of injected trout.

## Supplemental data file 1

**Differentially expressed genes in myogenic precursors from hyperplastic muscle *vs* non hyperplastic muscle.**

Heat map file for Java treeview visualisation of hierarchical clustering of differentially expressed genes in JT myogenic precursors from hyperplastic muscle *vs* non hyperplastic muscle (FJT and AT). (CDT 496 ko).

